# Timing of delamination of inner ear neurons specifies their topography and target innervation

**DOI:** 10.1101/2024.11.25.625134

**Authors:** Surjit Singh Saini, Raj K. Ladher

## Abstract

The neurons of the inner ear delaminate from a sensorineurogenic epithelium in the ventral part of the otocyst. Delaminated neuroblasts then condense to form the acoustico-vestibular ganglion (AVG). As they differentiate, the neurons connect mechanosensory hair cells (HCs) of the inner ear with their targets in the hindbrain in a precise topographical manner. However, it is unclear how or when positional identities of neurons are specified within the acoustico- vestibular ganglion (AVG), such that topographical information from HCs is maintained into the auditory centres of the brain. Here we find that the time of delamination from the otocyst correlates with neuroblast position in the ganglion. Using markers for neuronal differentiation, we find that the ganglion differentiates from a proximal to distal wave. Neurons that differentiate first innervate the vestibular apparatus, including the lagena and the proximal regions of the BP. Using sequential somatic cell labelling, we find that the central projection also follows a similar dependency on delamination order. Our studies show that the time of delamination of otic neuroblasts presage their target innervation choice and fibre positions within the developing eighth cranial nerve. We suggest that temporal information specifies the spatial identities during early inner ear neuron development.

## INTRODUCTION

Developing nervous systems must solve a non-trivial problem of neurons with positional identity, making the appropriate connections with the appropriate targets (Lumsden, 1993; Liu et al., 2001; Hochstim et al., 2008; Dessaud et al., 2010; Imai, 2012; Sagner and Briscoe, 2019). The vertebrate inner ear provides an excellent system to explore these questions. The complexity of this sensory organ contrasts with its initial development when it starts as a simple ectodermal thickening called the otic placode (Wu and Kelley, 2012; Sai and Ladher, 2015). All the sensory targets of the inner ear (eight in chick and six in mouse) arise from this simple placode after it has invaginated to form the otic vesicle (OV) (Wu and Oh, 1996).

Moreover, both vestibular and auditory neurons of the inner ear are placodal in origin. These arise from a restricted anteroventral domain of the OV called the pro-neurosensory domain (PNSD) (D’Amico-Martel and Noden, 1983; Hemond and Morest, 1991b). Initially, the precursors delaminate from the otic epithelium as neuroblasts and coalesce to form the acousticovestibular ganglion (AVG). From here, neuroblasts divide, differentiate, and then innervate the hair cells (HCs) of the inner ear, as well as neurons of the vestibular and auditory brainstem (Coate and Kelley, 2013). In the avian inner ear, another population of neurons innervate the lagena macula, a vestibular structure at the far end of the auditory organ that has been lost in placental mammals (Fischer et al., 1994; Kaiser and Manley, 1996; Khorevin, 2008). Concurrent with neurogenesis, the vestibular and auditory epithelium of the inner ear also develop (Fermin and Cohen, 1984; Hemond and Morest, 1991a).

The position of an inner ear neuron has implications on its fate (vestibular or auditory), tonotopic identity (for auditory neurons) and target innervation choice. However, how positional identities of the neurons of inner ear are determined needs to be clarified.

Experiments exploring the clonal relationships between neurons and HC, have ruled out a lineage relationship between target and nerve (Satoh and Fekete, 2005). Studies have shown that the tonotopic identities of HCs, which occupy different positions in the basilar papilla (BP) result from graded morphogen signalling (Mann et al., 2014; Thiede et al., 2014; Son et al., 2015). However, neuronal differentiation precedes that of HCs and thus how positional identity within the ganglion is established is unclear (Whitehead and Morest, 1985; Bok et al., 2013; Pyott et al., 2024). In the mouse, the AVG shows a gradient of differentiation (Ruben, 1967; Liu et al., 2010). This is reflected in the gradual loss of Shh expression in the ganglion, which is thought to control the onset of differentiation of post-mitotic progenitors into HC and SC in the auditory epithelium (Liu et al., 2010; Bok et al., 2013). This suggests that a pre-pattern does exist in the ganglion. By labelling Ngn1 positive neuronal progenitors at different times, it is apparent that the onset of neuroblast differentiation plays a role in the patterned innervation of both HC and central targets of ganglion neurons (Koundakjian et al., 2007). As neuroblasts are generated in the otic vesicle, and then undergo a limited number of divisions in the ganglion itself, it is unclear if the pattern within the ganglion is generated in the otic vesicle or through patterning cues acting on the ganglion once neuroblasts coalesce.

Studies in the chick suggest spatial patterning within the OV segregates vestibular and auditory neurons (Bell et al., 2008). Contrary data suggests that vestibular and auditory ganglion cells may not be strictly segregated in the chick inner ear (Satoh and Fekete, 2005; Hidalgo-Sanchez et al., 2023). In zebrafish, the timing of delamination has been shown to correlate with the position of neurons in the fish ganglion (Dyballa et al., 2017). Single-cell RNA sequencing in the mouse OV also revealed the dynamic nature of neuroblast delamination but did not reveal a particular pattern of heterogeneity amongst them (Matern et al., 2023). Thus, although temporal differences in neuroblast birth are critical in the formation of patterns in the ganglion, it remains unclear how and when these differences are encoded. Using tissue-specific labelling, we asked if temporal differences within the chick otic vesicle correlate with the position in the ganglion and, if so, how this influences the innervation pattern of inner ear epithelia. We find that the timing of delamination from the otic vesicle determines the position within the ganglion, with early delaminating neuroblasts occupying proximal positions and later delaminating occupying distal positions. We find that differentiation in the AVG follows a proximal to distal pattern. Early differentiating neurons innervate proximal targets in the BP. The exception is the lagenar neurons that are born early but innervate HC at the distal end of the BP. Topographic targeting shown as a consequence of neuroblast delamination and differentiation is preserved in the central projections, suggesting that the temporal control of delamination within the otic vesicle is essential for the initial peripheral and central topographic maps of inner ear innervation.

## METHODS

### Chick Embryos

All protocols involving the use of fertilised chicken eggs and unhatched embryos followed institutional guidelines and were approved by the Institutional Animal Ethics Committee of the National Centre for Biological Sciences, Bengaluru, Karnataka. Fertilised eggs were obtained from the Central Poultry Development Organisation and Training Institute, Hasserghatta, India. Eggs were incubated in a humidified chamber at 38°C with humidity maintained at 45% for the appropriate length of incubation. They were staged, according to Hamburger and Hamilton (Hamburger and Hamilton, 1992).

### Sectioning

Embryos were fixed in 4% paraformaldehyde (PFA) at 4⁰C overnight, washed in PBS and cryoprotected by passing through a sucrose series and embedded in OCT medium (UF1000 Cancer Diagnostics). Heads were sectioned coronally at a thickness of 30 µM on charged slides and dried at 37⁰C overnight.

### Immunohistochemistry

Immunohistochemistry was performed as previously described (Singh et al., 2022). Briefly, sections were washed in PBS and permeabilised with 0.5% Triton X-100 (T8787, Sigma-Aldrich), followed by blocking with PBS + 0.5% Triton + 1% BSA + 3% goat serum.

Primary antibodies used were Tuj1 (T2200, Sigma-Aldrich, 1:500), NeuN (MAB377, Merck Millipore, 1:300), GFP (G6539, Sigma-Aldrich,1:500), mCherry (PA5-34974, Thermo Fisher, 1:500), Neurofilament (3A10C, DSHB, 1:300), Laminin (PA1-16730, Thermo Fisher, 1:500) and Myosin 7A (Myo7A 138-1, DSHB, 1:300). Secondary antibodies used were (A11037, goat anti-rabbit, Alexa fluor 594, Invitrogen, 1:500), (A11034, goat anti-rabbit, Alexa fluor 488,invitrogen, 1:500), (A11005, goat anti-mouse, Alexa fluor 594, Invitrogen, 1:500), (A11029, goat anti-mouse, Alexa fluor 488, Invitrogen, 1:500), (A21203, donkey anti-mouse, Alexa fluor 594, Invitrogen, 1:500) and (A32733, goat anti-rabbit, Alexa fluor 647,Invitrogen, 1:500). DAPI (D1306, Thermo Fisher, 1:2000) was used as a nuclear stain.

### EdU Click Reaction

Fertilized eggs at the desired stage were windowed, and the vitelline membrane was removed carefully only around the head region. EdU (Invitrogen C10337) was diluted in 0.719% filtered saline, with the concentration and volume of EdU optimized for different staged embryos. For embryos grown to HH 26 (E5) stage, 100 µL of 100 µM EdU was added at HH 17/18 (E3) or HH 23 (E4). For embryos grown till HH 29 (E6) stage, 200 µL of 150 µM EdU was added at stage HH 17/18 (E3), HH 23 (E4) or HH 26 (E5) stage. For EdU labelling at HH 29 (E6) stage 600 µL of 150 µM solution was added and embryos incubated only for 10 to12 hours. In all cases, half of the solution was pipetted directly onto the OV and remaining solution over the entire embryo. Eggs were sealed and re-incubated until the desired stage, and then fixed in 4% PFA overnight at 4⁰C, and then sectioned. The click reaction was performed as manufacturer’s protocol (Invitrogen C10337), followed by immunostaining with appropriate antibodies.

### EdU and BrdU dual labelling

Dual labelling of proliferating neurons was possible only for early stages of chick embryos as the lethality was high when embryos were subjected to both the reagents - Edu and BrdU. 100 µL of 100 µM EdU (Invitrogen C10337) was added to the embryos at stage HH 18 and embryos were incubated till stage HH 22. At stage HH 22, BrdU (Invitrogen 000103) was diluted from the stock solution in 0.719% filtered saline to a ratio of 1:10 and the same was pipetted over the embryos. Embryos were cultured to stage HH 23 and fixed in 4% PFA overnight at 4⁰C. The next day, PFA was removed and embryos were washed in PBS multiple times. Cryosections were obtained as mentioned in immunohistochemistry (section 1). EdU click reaction was performed as mentioned above (in section 2). After the click reaction, slides were washed in PBS and briefly dipped in dilute HCl in Coplin jars at room temperature for around 20 to 30 minutes. This was done for antigen retrieval before proceeding towards BrdU immunostaining. The Coplin jar was wrapped in aluminium foil, and the slides were light-protected during the entire process. Slides were removed and washed multiple times on a shaker by dipping in a fresh Coplin jar containing PBS. In order to ensure no cross-reactivity between EdU and BrdU during immunostaining, a mouse monoclonal antibody (Clone MoBU-1, Invitrogen B35132), which has been verified to be specific to BrdU, was used. Sections were permeabilized with 0.5% Triton X-100. This was followed by a blocking solution (1% BSA + 3% goat serum) for 1 to 2 hours. BrdU antibody (Invitrogen B35132) and Tuj1 antibody (Invitrogen) diluted in blocking solution at concentrations of 1:300 and 1:500 respectively was added and sections were incubated overnight at 4⁰C. The following day after brief washing with PBS, secondary antibodies were added. Alexa fluor 594 conjugated secondary antibody was used for BrdU detection (donkey anti-mouse, Invitrogen, A21203, Alexa fluor 594).

### Electroporation

#### Single electroporation

After reaching the desired stage, eggs were windowed by opening a part of the shell. The vitelline membrane surrounding the embryonic head was removed to access the OV. The OV cavity was carefully filled with a pulled glass needle with a mixture of plasmid, sucrose (for increasing viscosity) and fast green (for visualisation). The final concentration of pCAG-eGFP or pCAG-mcherry was 1 µg/µL. pCAG-eGFP was a gift from Wilson Wong (Addgene plasmid # 89684 ; http://n2t.net/addgene:89684 ; RRID:Addgene_89684).

CAG-mCherry was a gift from Jordan Green (Addgene plasmid # 108685 ; http://n2t.net/addgene:108685 ; RRID:Addgene_108685). Only the OV cavity was filled, and care was taken to ensure that neurons and surrounding mesenchyme were not injected directly. As a result, only neurons which delaminate at or after the injection time take up the electroporated plasmid. The positive electrode was inserted below the embryo via an opening on the blunt side of the egg. The negative electrode was held over the injected otocyst.

Electric pulses were given by using a foot pedal of CUY21 electroporator (NEPA gene). Voltage was optimized for different stages (12 V for HH 17, 18 V for HH 22). 5 pulses of current separated by 50 milliseconds were used. 100 to 200 µL of filtered 0.719% saline was added on top of the embryo after electroporation, and the egg resealed with scotch tape and placed back for incubation.

#### Sequential electroporations

Here the same OV of an embryo was electroporated at two different time points (separated by 24 hours). The first electroporation is done at HH 17 stage with pCAG-mcherry plasmid and the second with a pCAG-GFP plasmid at HH 22 stage. The window of the egg after its first electroporation was reopened on the next day. Usually, the vitelline membrane grows back over the exposed OV and has to be removed. The HH22 otocyst was carefully filled, ensuring that only the cavity is filled. The positive electrode was reintroduced from the blunt end. We note that embryo survival is reduced if albumin leaks from this opening. The cathode was placed above the otocyst and electroporated. Filtered 0.719% saline was added onto the embryo. Embryos were grown for two more days and fixed at HH 29 (E6) stage.

#### Sparse labelling of delaminating neurons

CREMESCLE was used for sparse labelling of delaminating cells (Schohl et al., 2020). We used a ratio of 1:10000 pCAG-Cre:pCALNL-GFP plasmids. Plasmids were obtained from Addgene: pCAG-Cre (Addgene 13775) and pCALNL-GFP (Addgene 13770). OV was electroporated with a mixture of pCAG-Cre, pCALNL-GFP, sucrose and fast green either at stage HH 17 or HH 22. The embryos were grown till stage HH 29 (E6).

### Imaging and analysis

Images were taken using FV3000 confocal inverted microscope. Image J was used for counting of cells and analysis. The inner ear regions were considered innervated if neurites were seen crossing the laminin barrier surrounding it. All statistical analysis was done using GraphPad Prism 10 software.

To define proximal and distal regions of the ganglion, a straight line was drawn at the long axis of the ganglion. At the midpoint of this line, the tissue was bisected in two halves. The part closer to the saccule and the proximal BP was called proximal ganglion. The other half which is away from saccule and towards the distal BP was called distal ganglion.

The same method was used to define the regions of BP that are innervated. A straight line was drawn along the long axis of BP, and its midpoint was chosen to define the proximal and distal regions. Innervation was said to be proximal if the neurites projected to the left of this midpoint and distal if seen on the right side.

## RESULTS

### Differentiation of chick inner ear neurons is patterned and follows a proximal to distal wave

To investigate the pattern of neuronal differentiation in the AVG, we stained the ganglia with a marker for differentiated neurons, neuronal nuclei (NeuN). The ganglia were co-stained with the neuronal marker Tuj1. Neuroblasts have started †o delamination from the OV by HH 19 (Fig 1A) and by HH 29 have coalesced to form the AVG (Fig. 1B). Plane of sectioning of embryos which was kept uniform for all experiments (Fig. 1C). 1D shows the orientation of axes.

**Figure 1.**
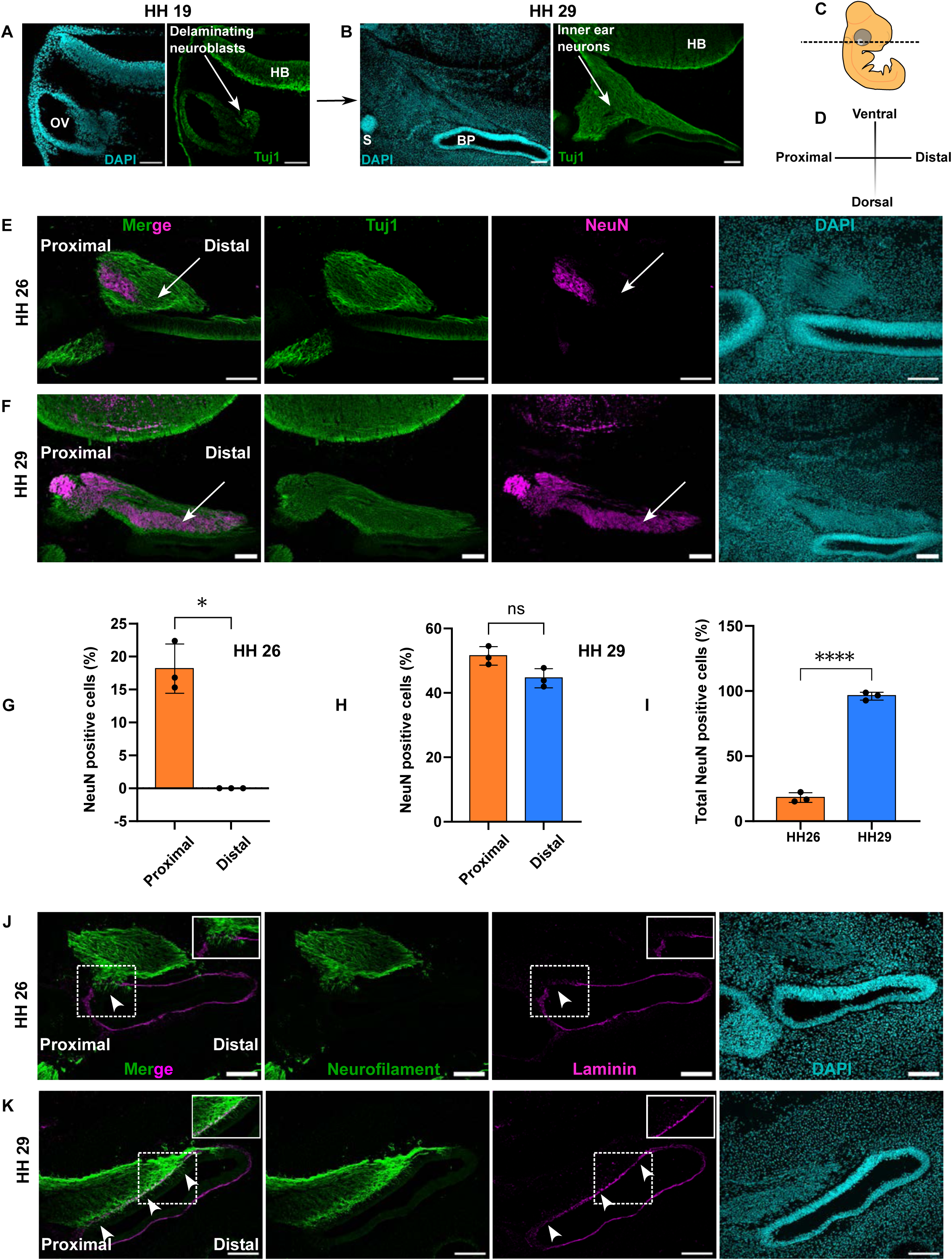
Patterned differentiation of chick inner ear neurons. A - Section from stage HH 19 chicken embryo showing otic vesicle (OV) and delaminating neuroblasts. B - Stage HH 29 section showing the acoustic-vestibular ganglion. (BP - basilar papilla, S - saccule, HB - hindbrain) C - Dotted line through the chick embryo shows the plane of sectioning followed in all experiments to get a proximal-distal orientation of inner ear and its ganglion. D - Axes showing the orientation of proximal-distal and dorsal-ventral regions in the images. E - F are sections from HH 26 (E) and HH 29 (F) embryos respectively. Proximal (left side) and distal (right side) regions have been kept uniform across all figures and the same has been shown. Arrows in E points to distal regions of the ganglion at stage HH 26, which are devoid of NeuN staining. Arrows in F point distal regions of the ganglion at stage HH 29, which at these stages show NeuN expression. G - Number of NeuN positive cells in the proximal and distal regions of HH 26 inner ear ganglion. Paired t test, N = 3 embryos, P value = 0.0138 (*) H - Comparison of NeuN positive cells in the proximal and distal regions of HH 29 inner ear ganglion. Paired t test, N = 3 embryos, P value = 0.1383 (ns - not significant) I - Comparison of total NeuN positive cells in the inner ear ganglion at HH 26 and HH 29 stage. Unpaired t test, sample size N = 3 embryos, P value < 0.0001 (****). J and K - Staining to show the spatio-temporal profile of innervation of the BP. J - At stage HH 26 neurofilament fibres are first seen ingressing into proximal regions of the BP (arrow head). Laminin breaches can be observed at the region of entry of these fibres (arrow head). K - At stage HH 29 the entire region (proximal, middle and distal) of the BP has been innervated by neuronal fibres (arrow heads). Concurrently breaches are now visible along the entire laminin expressing region (on the neural side of BP). Scale bar 100 μm

We failed to detect appreciable expression of NeuN at stages earlier than HH 26 (E4.5-5). At HH 26, we observed NeuN in the proximal-most regions of the ganglion (arrows in Fig.1E show the absence of NeuN in distal regions). As development proceeded the expression extended progressively more distal (arrows in Fig.1F). By stage HH 29 the entire ganglion was positive for NeuN expression (Fig. 1F, I). This indicated that all inner ear neurons had become post-mitotic and had differentiated by stage HH 29. Moreover, patterned differentiation occurs in chick AVG following a proximo-distal direction. This was can be seen even in the ventral sections of the ganglion (Supplementary Fig.1S A and B).

Classical studies in chick have studied the innervation patterns during early inner ear development. These report breaches in the basal lamina of the otocyst as indicative of the time of neurite entry (Hemond and Morest, 1991a). To further corroborate the proximal to distal wave of differentiation of AVG neurons, we used this criterion for assessing innervation. We reasoned that the innervation pattern of the ear may parallel the differentiation wave observed in the AVG. As the neurites invade the sensory epithelium, local breaches in the lamina surrounding those regions would be seen. We used neurofilament immunostaining to label neuronal fibres and an antibody against laminin to label the

extracellular basal lamina surrounding the nascent BP. The developing BP showed intact laminin expression till stage HH 24/25 (see Supplementary Fig.1SC) and no ingressing neurites into the BP. At stage HH 26, when the first NeuN expression was seen in the proximal regions of the ganglion, we observed innervation of the proximal-most part of the BP and breaches in the laminin in that region (arrowheads in Fig.1J and inset). By stage HH 29, laminin breaches and invasion of neurites was detected along the entire proximal-distal axis of the BP (arrow heads in Fig.1K. Inset shows entry of neurites even in the distal regions of BP at HH 29). Thus, like differentiation, the innervation wave also proceeds following a proximal to distal direction in the developing inner ear epithelia.

### Birth timing of inner ear neurons specifies their positions in the ganglion

Our data indicates that neuronal differentiation in the AVG proceeded as a proximo-distal wave. We next asked if the proliferation order also followed a similar pattern. As the positional identity of neurons in cortical layers is known to depend on the timing of their birth (McConnell and Kaznowski, 1991; Campbell, 2005) we wondered if positions of neurons of chick inner ear ganglion could also be related to their birth-order. Using an in ovo EdU labelling protocol (Fig.2A), we marked cells in the S-phase of the cell cycle at different developmental times. EdU was added to the embryos either at stage HH 19/20 or HH 23 and incubated until HH 26. At this stage, embryos were fixed, sectioned, and stained for NeuN (which is restricted to the proximal ganglion at HH 26) and EdU reactivity revealed. We observed that EdU application of HH 19 resulted in labelling in the proximal parts of the AVG (Fig. 2B). Here, the proximal expression of NeuN in the ganglion could be seen at the same place (arrowhead in Fig. 2B) as these early born neurons. Moreover, these early born neurons were absent from distal regions of the ganglion (arrow in Fig. 2B). In contrast, the HH 23 application of EdU resulted in neurons occupying distal ganglion positions (Fig. 2C).

**Figure 2.**
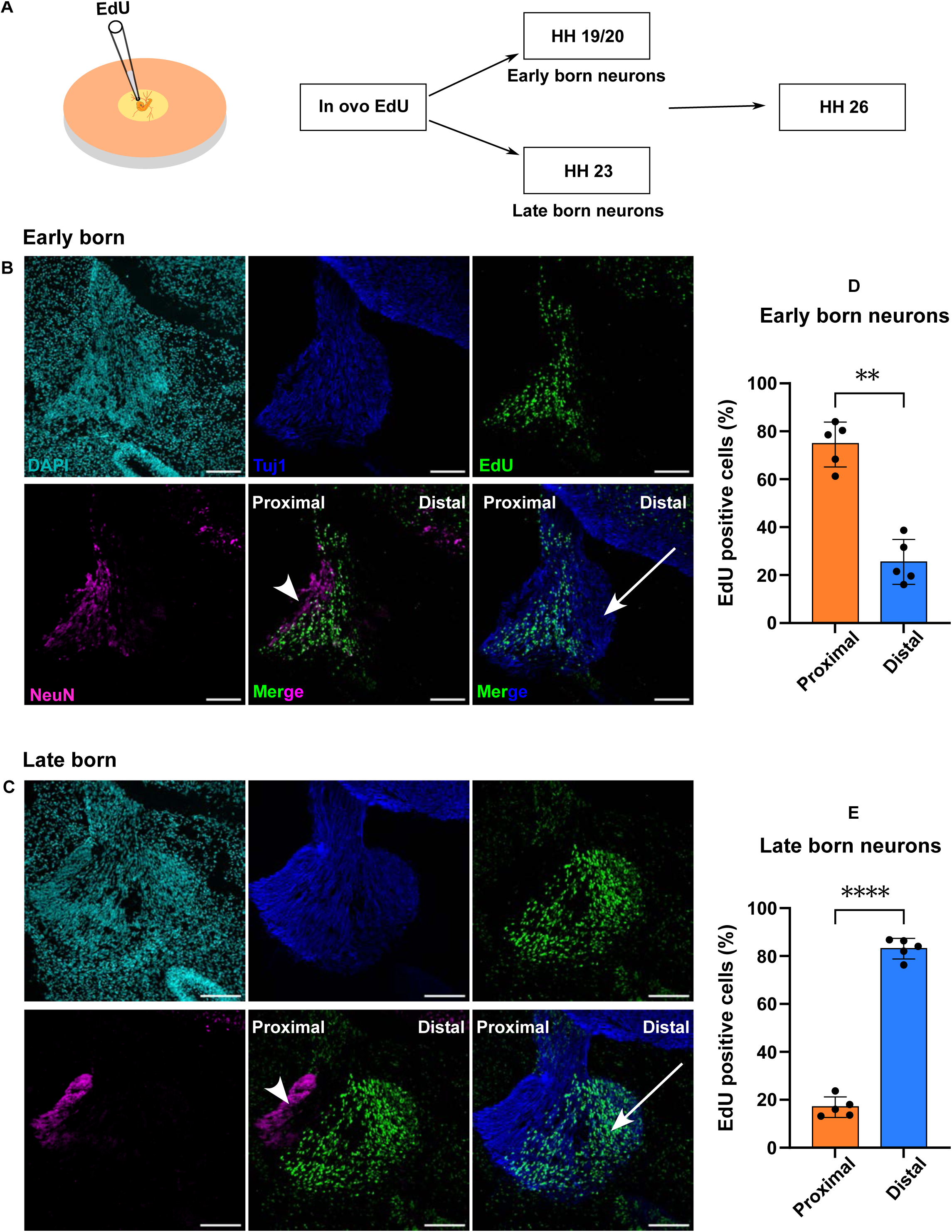
Birth timing and position of neurons at HH 26 stage. A - Experimental protocol for labelling proliferating cells at different developmental times. In ovo EdU at stage HH 19/20 labels early-born neurons, while at HH 23 labels late-born neurons. B - Sections showing early born neurons. NeuN expression which at stage HH 26 is restricted to proximally was used as a regional marker. Arrowheads show the proximal position of early born EdU positive neurons with respect to NeuN expression. Arrow point to distal regions of the ganglion, from where early born neurons are absent. C - Sections showing late-born neurons. Arrowhead points to the proximal region (marked by NeuN), which is devoid of late-born EdU positive neurons. Arrow point to distal regions of the ganglion and show late-born neurons in these regions. D - Comparison between proximal and distal positions of early born neurons within the same ganglion. Paired t test, N = 5 embryos, P value = 0.0043 (**) E - Comparison between proximal and distal positions of late-born neurons within the same ganglion. Paired t test, N = 5 embryos, P value = 0.0001 (****) Scale bar 100 μm

The NeuN expressing region was distinctly separated from regions occupied by these late born neurons (arrowhead in Fig. 2C). We next asked if sequential labelling, first with EdU and then with BrdU could segregate these populations of neurons in the same sample (Supplementary Fig. 2SA). As lethality was high once BrdU is applied after EdU, we incubated these samples to HH 23. Similar to the single labelling, we observed EdU positive early-born neurons were proximal. In contrast, BrdU positive late-born neurons were found at distal locations in the ganglion (see arrowheads in Supplementary Fig.2SE). Thus, our data suggests early and late proliferating neurons occupy distinct proximo-distal locations in the AVG.

### Inner ear neurons become post-mitotic in a proximal to distal manner

Our NeuN expression observations point to a proximal to distal wave of differentiation in the chick AVG. To better understand this, we performed in ovo EdU labelling at stages HH 19/20, HH 23, HH 26 and HH 29, and incubated the embryos to stage HH 29 (except embryos that received EdU at HH 29, which were fixed at stage HH 30) (Fig. 3A).

**Figure 3.**
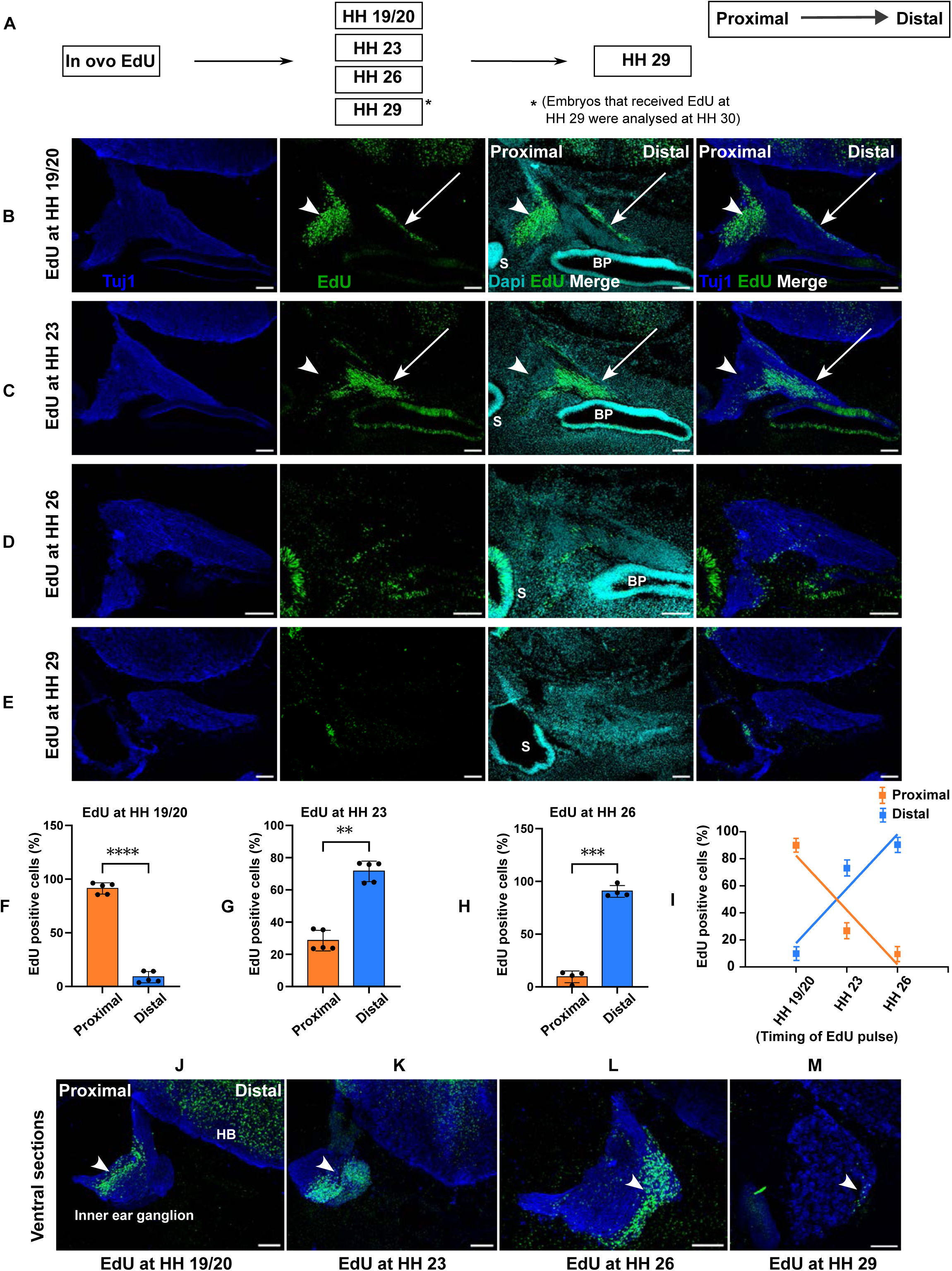
Earliest-Born Neurons show proximal and distal locations in the AVG. A - Protocol used for labelling of proliferating cells at different developmental stages. B - Sections showing inner ear neurons from an embryo that received EdU at HH 19/20. Arrow heads show EdU positive cells in the proximal region of the ganglion. Arrow shows a small group of EdU positive cells found flanking the distal regions of the ganglion. C - Sections showing inner ear neurons from an embryo that received EdU at HH 23. Arrow heads show absence of EdU positive cells in the proximal regions of the ganglion (cf. corresponding regions in B). Arrows show the flanking regions of distal ganglion devoid of EdU staining (cf. corresponding regions in B) D and E are sections from embryos that received EdU at HH 26 (D) and HH 29 (E) respectively. (S- saccule, BP- basilar papilla) F- Comparison of proximal and distal positions of proliferating inner ear neurons in embryos that received EdU at HH 19/20. Paired t test, N = 5 embryos, P value < 0.0001 (****) G - Comparison of proximal and distal positions of proliferating inner ear neurons in embryos that received EdU at HH 23. Paired t test, N = 5 embryos, P value < 0.0017 (**) H - Comparison of proximal and distal positions of proliferating inner ear neurons in embryos that received EdU at HH 26. Paired t test, N = 4 embryos, P value < 0.0007 (***) I - Neuronal birth timing and their position in the ganglion across developmental stages. Simple linear regression, N = 4 embryos for each developmental stage, R squared = 0.8852 and P value < 0.0001 (proximal) and, R squared = 0.8852 and P value < 0.0001 (distal) J - M are ventral sections from embryos that received EdU at HH 19/20 (J), HH 23 (K), HH 26 (L) and HH 29 (M). Arrowheads show the proliferation wave gradually shifting along the proximo-distal axis of the ganglion. Very few EdU positive cells (< 10) are detectable in the inner ear ganglion of embryos that received EdU at HH 29 stage. (arrowhead in M). Embryos in B, C and D were all analysed at HH 29. However, embryos that received EdU at stage HH 29 (E) were analysed at HH 30. Scale bar 100 μm

The proximal expression of NeuN at HH 26 allowed us to unequivocally define position, but even at these stages, the ganglion is undergoing development. As well as proliferation, the ganglion undergoes morphogenesis after HH 26, such that it elongates along the proximal-distal axis. Importantly, the separation of the vestibular and auditory components of the AVG become distinct. As discussed already, and similar to the observations at HH 26, at HH 29 cells labelled with EdU at HH 19/20 are predominantly localised to the proximal parts of the AVG, with contributions to both vestibular and auditory regions (arrowheads in Fig.3B). However, we observed EdU positive neurons in the border regions of the distal ganglion (arrows in Fig.3B). We reasoned that as the ganglia underwent morphogenesis, some HH 19 EdU-labelled neurons migrated to these distal regions. These neurons are temporally related to vestibular neurons, we speculated that these are lagenar neurons. HH 23 EdU-labelled neurons were found in the distal ganglion and not present in the proximal-most parts (arrowheads in Fig. 3C). These were also distinct from the regions that we identified as being occupied by lagenar neurons (arrows in Fig. 3C). Thus, our repetition of EdU labelling of different stages of ganglion and incubation to stage HH 29, allowed us to make fortuitous observations on the birth timing of lagenar neurons which to our knowledge was not known previously. Despite being spatially segregated after stage HH 26 from other vestibular neurons, we found they were specified at the same time as their proximal vestibular counterparts.

By stage HH 26 proliferation in the ganglion had reduced and ceased by HH 29 (Fig. 3D and E). When ventral regions of inner ear ganglion from embryos EdU labelled at stage HH 19/20, HH 23, HH 26 and HH 29 were analysed, a clear wave of proliferation progressing from proximal to increasingly distal regions was observed across the ganglion (arrow heads in Fig. 3J, K, L and M). This implies that just as in mammalian inner ear ganglion where cell cycle exit (CCE) proceeds from base to apex, the avian inner ear neurons also undergo cessation of proliferation with conserved directionality from proximal to distal regions.

### Timing of delamination of otic neuroblasts specifies the positions of neurons in the inner ear ganglion

Neurons of the AVG are generated from neuroblasts that leave the otic epithelium through delamination. This process occurs over a protracted period of around two days in the chick from around HH 12/13 to HH 22/23- (Alvarez et al., 1989; Hemond and Morest, 1991b). We were intrigued by the possibility that these temporal delamination differences may affect the positional identities of inner ear neurons. Using in-ovo electroporation, we targeted a plasmid constitutively expressing eGFP (pCAG-GFP) to the otic vesicle at two different developmental time points, HH 18 and HH 22/23- (Fig. 4A). We studied the positions of GFP-labelled delaminating neurons at stage HH 26, using NeuN as a proximal marker of the ganglion. Subsequent delamination experiments were performed to study the ganglion at stage HH 29, where the proximal-distal axis is anatomically more pronounced.

**Figure 4.**
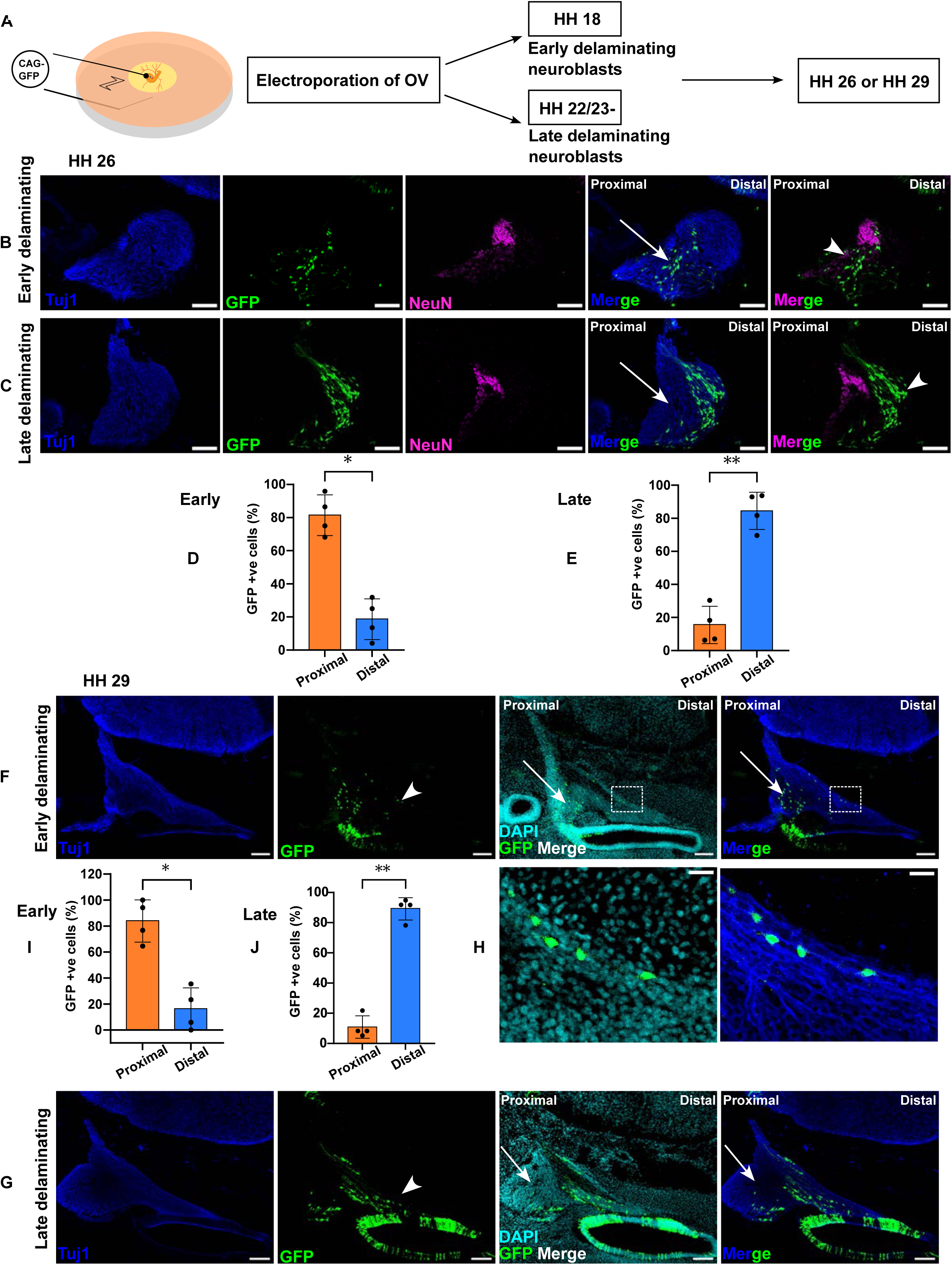
Timing of delamination and position of inner ear neurons. A - Schematic representation of electroporation protocol for labelling early (HH 18) or late (HH 22/23-) delaminating neuroblasts. B - Sections of stage HH 26 embryos showing early delaminating neuroblasts. C - Sections of stage HH 26 embryos showing late delaminating neuroblasts. Arrowheads show position of GFP labelled delaminated cells with respect to NeuN expression. D - Comparison of positions occupied by early delaminating neurons in the ganglion at stage HH 26. Paired t test, N = 4 embryos, P value = 0.0145 (*) E - Comparison of positions occupied by late delaminating neurons in the ganglion at stage HH 26. Paired t test, N = 4 embryos, P value = 0.0089 (**) F - Sections of stage HH 29 embryos showing early delaminating neuroblasts. G - Sections of stage HH 29 embryos showing late delaminating neuroblasts. H - Zoomed image of the boxed region in F. I - Comparison of positions occupied by early delaminating neurons in the ganglion at stage HH 29. Paired t test, N = 4 embryos, P value = 0.0245 (*) J - Comparison of positions occupied by late delaminating neurons in the ganglion at stage HH 29. Paired t test, N = 4 embryos, P value = 0.0018 (**) Scale bar 100 μm in B, C, F and G and 20 μm in H

At stage HH 26, early delaminating cells were found only in the proximal, NeuN positive region of the ganglion (arrowhead in Fig. 4B). Late delaminating cells were excluded from proximal positions (arrow in Fig. 4C) and instead inhabited the distal regions of the ganglion (arrowhead in Fig. 4C). This suggested that the positional identities of inner ear neurons were specified at the time of precursor delamination from the OV, that temporal differences in delamination order specify a positional order. In HH 29 ganglia, early delaminated GFP-positive cells were seen in the proximal region (arrows in Fig. 4F), and a few delaminated cells could be seen in the distally located lagenar ganglion (arrowhead and boxed region in Fig. 4F; the boxed region has been zoomed and shown in H). Late delaminating GFP-positive cells were confined to the distal areas of the auditory ganglion at HH 29. They were absent from the proximal regions of the ganglion (arrows in Fig. 4G).

Taken together with the results of birth timing of inner ear neurons, the delamination timing results imply that cells that leave the OV earlier proliferate first and take up proximal identities, while cells that delaminate at later time points proliferate late and become distal.

### The timing of otic neuroblast delamination positions central projections in the 8^th^ nerve

The ganglion neurons are bipolar, and as well as projecting to the auditory epithelium, send projections to the centre. As tonotopy is maintained in the central projections, we asked whether the order imposed by the timing of delamination of otic neuroblasts, perdured in the central projections. To test this, we used a sequential electroporation protocol. The OV was electroporated with a pCAG-mCherry expressing plasmid at HH 18, reincubated for a day, and then electroporated with a pCAG-eGFP plasmid.The embryos then incubated to HH 29 (Fig.5A).

**Figure 5.**
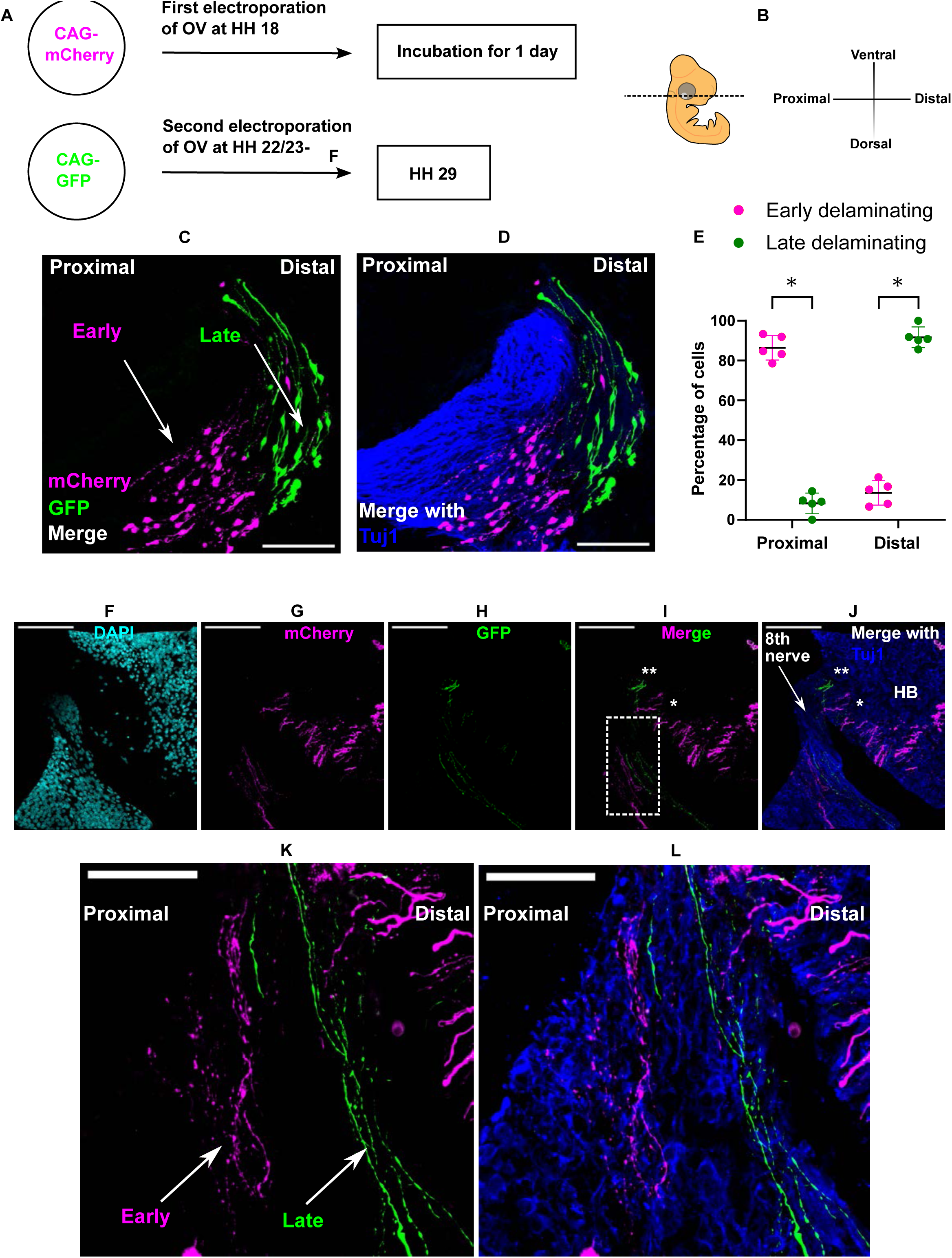
Timing of delamination of otic neuroblasts and topography of the 8^th^ nerve. A - Protocol showing sequential labelling of early (HH 18) and late (HH 22/23-) delaminating neuroblasts by successive electroporation of same OV at different developmental times. The embryos were incubated to stage HH 29. Position of neurons in the ganglion and those of fibres from early (mCherry positive) and late (GFP positive) delaminating neuroblasts were analyzed in the 8^th^ nerve. B - Sectioning plane and orientation of axes. C - Section showing merge of mCherry and GFP positive neurons. D - Section showing staining of Tuj1, mCherry and GFP merge. E - Comparison of time of delamination of cells with their position (proximal or distal) in the same ganglion. Multiple paired t test, N = 5 embryos, P value = 0.000007 (*) for proximal, P value = 0.000007 (*) for distal. Scale bar 100 μm in C and D F - Stage HH 29 DAPI sections at the level of nerve entry into the hindbrain. G - mCherry positive fibres from early delaminating neurons. H - GFP positive fibres from late delaminating neurons. I - Merge of mCherry (early) and GFP (late) fibres. J - Merge of Tuj1, mCherry and GFP. Early fibres are seen more caudal (*) and late fibres more rostral (**) on their entry into the hindbrain (in I and J). K and L - Magnified image of the boxed region in I. K - mCherry and GFP merge image showing spatial segregation of fibres from early and late delaminating neurons. L - Tuj1, mCherry and GFP merge. Scale bar 100 μm in F, G, H, I and J and 50 μm in K and L

In agreement with our single electroporations, we observe that early delaminating, mCherry positive neurons are found in proximal parts of the ganglion, and later delaminating, eGFP positive neurons are found in the distal part of the ganglion (arrows in Fig.5C). In the central tracts, we observed segregation of mCherry from eGFP labelled fibres (arrows in Fig. 5K and L). An earlier study which had used DiI and DiD labelling of proximal and distal regions of the chick BP and cochlear ganglion, had shown that neurons that labelled these tonotopically distinct regions of the auditory epithelium have fibres that are topographically arranged in the cochlear nerve (Molea and Rubel, 2003). Our protocol labels the precursors that just delaminate from the OV (much before the appearance of any targets). Despite the difference in labelling paradigm, we find that the axons of neurons that delaminate early or late and innervate tonotopically distinct regions of the BP, are topographically segregated in the 8^th^ nerve. The orderly arrangement of early and late delaminating fibres in the nerve from precursors that leave the otic epithelium at different times is suggestive that the information of nerve topography may very well lie (at least partially) in the temporal order of delamination. Although we did not trace these fibres towards the vestibular or auditory nucleus as we restricted ourselves to stage HH29, we did notice that the early mCherry labelled fibres are seen more caudal (* in Fig. 5I and J) while the late GFP labelled fibres are more rostral in the developing hindbrain (** in Fig. 5I and J).

### Timing of delamination specifies target innervation choice

To understand if the timing of delamination also correlated with the position of target innervation, we employed a Cre-Lox based protocol, CREMSCLE, to sparsely label delaminating neuroblasts (Schohl et al., 2020). Here, a higher concentration of the pCALNL- eGFP expressing plasmid was co-electroporated with pCAG-Cre expressing plasmid at a 1/8000 fold lower concentration into the OV, at either HH 18 or HH 22/23- (Fig. 6A).

**Figure 6.**
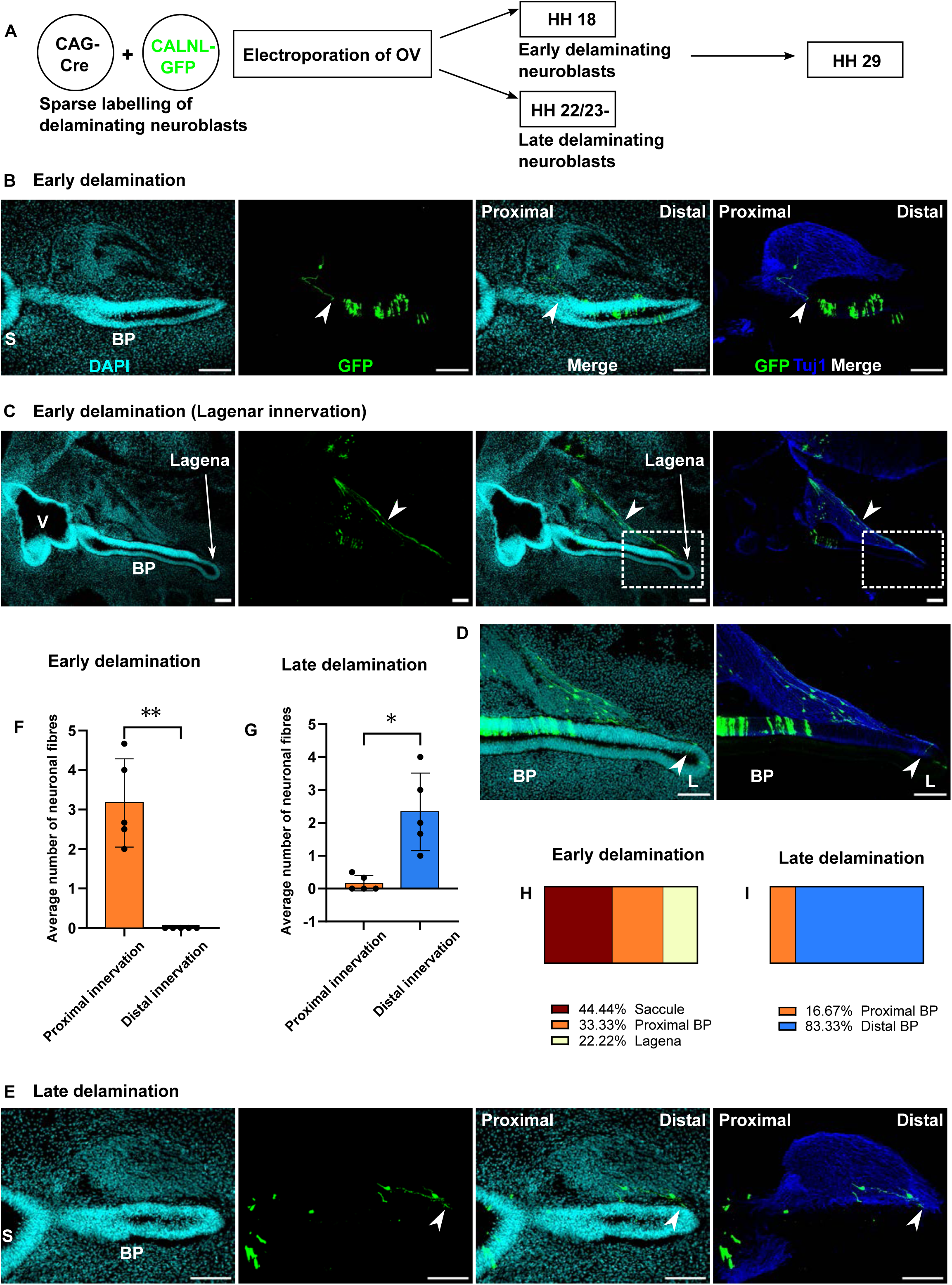
Timing of delamination and target specification. A - Protocol used for sparse labelling of delaminating neuroblasts from the OV. A mixture of plasmids pCALNL-EGFP (at high concentration) and pCAG-Cre (at very low concentration) was co-electroporated into the OV to sparsely label delaminating neuroblasts at stages HH 18 (early delaminating) and HH 22/23- (late delaminating). Their target epithelium was analyzed at stage HH 29. B - Panel showing sparse labelling of early delaminating neurons. Arrow head shows the target region - proximal part of BP of an early delaminated GFP positive cell. (S- Saccule, BP- Basilar papilla) C - Panel showing sparse labelling of early delaminating neurons (which contribute to the lagenar ganglion, arrow head) innervating the lagena macula (boxed region). D - Zoomed image of the boxed region in D to show the innervation of lagena (arrow heads). E - Panel showing sparse labelling of late delaminating neurons. Arrow head shows the target region, the distal part of BP of late delaminated GFP positive cells. F - Comparison of different targets innervated by early delaminating neurons. Paired t test, N = 5 embryos, P value = 0.0032 (**) G - Comparison of different targets innervated by late delaminating neurons. Paired t test, N = 5 embryos, P value = 0.0193 (*) H - Horizontal slice showing relative contribution to the innervation of different target regions of a single inner ear by early delaminating neurons. I - Horizontal slice showing relative contribution to the innervation of different target regions of a single inner ear by late delaminating neurons. Scale bar 100 μm

pCAG-mcherry was also electroporated into the OV as a tracer, both to verify the extent of electroporation and to ensure OV targeting only. Embryos were incubated to HH 29. Sparse nature of labelling was first validated (see dotted circle in supplementary Fig.6SA- c, d and f).

We find that early delaminating neuroblasts populate the proximal part of the ganglion, and innervated the proximal region of the BP (arrowhead in Fig. 6B). Early delaminating neurons were also seen innervating the saccule, which is adjacent to the proximal BP (arrows in supplementary Fig. 6SB- d and e). As the vestibular hair cells have differentiated at HH 29, we immunostained saccular sections with a Myo7a antibody showing this innervation of vestibular HCs. (see supplementary Fig. 6SC-b and c. Arrows show a fibre innervating utricle and arrowheads show innervation of HCs in the saccule). This implies that early differentiating HCs are innervated by neurons that delaminate earlier from the wall of OV. We also observed early delaminating cells innervating the lagena, with highly stereotyped projections at the edge of the auditory ganglion, turning into the developing lagena macula (Fig. 6C and arrowheads in Fig. 6D). We did not observe any misrouting of the lagenar fibres into the BP via the auditory ganglion. Otic neuroblasts, electroporated at HH 22/23-, populated distal parts of the ganglion and innervated more distal regions of the BP (arrowheads in Fig. 6E). We were unable to detect any vestibular or lagenar innervation by these late delaminating neuroblasts.

## DISCUSSION

Neuronal birth order information can establish positional identity and topography across various developing systems. The dependence on birth order has been observed in other developing neural systems. Neurons born at different times occupy specific laminar positions during corticogenesis (McConnell and Kaznowski, 1991; McConnell, 1992). Studies focusing on the auditory portion of the AVG have also highlighted the link between birth order and pattern. Work in the mouse inner ear had investigated the birth timing of neurons within the ganglion, revealing cell cycle exit (CCE) from base (proximal) to apex (distal), a pattern that we find is conserved in birds (Ruben, 1967). The P-D CCE direction corresponds to the P-D order of innervation of hair cells within the organ of Corti(Koundakjian et al., 2007). Furthermore, this is reflected in topographical projections to the central targets of cochlear nucleus (Scheffel et al., 2020). It is also known that neurons of the mouse anteroventral cochlear nucleus (AVCN) are arranged according to their birthdates, which is aligned with the tonotopic axis (Shepard et al., 2019). In this study, we focused on the production of the neuroblasts within the otocyst. We found that the downstream events of birth order, positional identities and even innervation choice are already specified in the otocyst encoded temporally when neuroblast progenitors delaminate.

### Innervation of inner ear epithelia

Neurites from the AVG reach both peripheral and central targets in a topographical fashion before any overt sign of differentiation (Hemond and Morest, 1991a; Molea and Rubel, 2003; Koundakjian et al., 2007; Scheffel et al., 2020). This implies a pre-pattern already exists in the inner ear ganglion. This prepattern is not only apparent in the choice of target but also in the pattern of cell cycle exit and subsequent differentiation. One of the clearest distinctions is that between vestibular and auditory neurons. Previous work in the chick had suggested that vestibular and auditory neurons were positionally segregated in the otocyst (Bell et al., 2008). However, more recent work, using a chimeric quail-chick model failed to find any clear spatial segregation of vestibular and auditory neuronal precursors in the otic placode (Hidalgo-Sanchez et al., 2023). The segregation is likely temporal, and work in the mouse, using an inducible neuroblast-specific Cre (Ngn1-CreERT2), found that only vestibular neurons were labelled when induction was performed early (Koundakjian et al., 2007).

Similarly, our studies find that early-born neurons labelled with EdU at HH 19 project to the vestibular sensory epithelia of the saccule and the lagena. The saccule and lagena flank the auditory epithelium proximally and distally, respectively. The observation that the lagenar neurons follow a stereotypical migration around the auditory neurons underscores the specification of early-delaminating neurons into vestibular neurons, and that the temporal progression of neurogenesis does not simply lead to progressively more distal targets being innervated.

Our sequential electroporation experiments indicate that the birth dating of inner ear neuroblasts also influences their central projections. This is consistent with data involving the unilateral removal of an otocyst from the developing chick embryo. This revealed that the central tonotopic pattern in the chick cochlear nucleus was indistinguishable from the contralateral unoperated side (Lippe et al., 1992). This implies that the initial tonotopic organisation of central targets is independent of peripheral inputs.

### Tonotopic order of auditory neurons

Our data shows that the positional identities of inner ear neurons are established at the time of their delamination. As the position of auditory neurons correlate to their eventual frequency responsiveness, it is natural to ask when is tonotopic identity specified? We have shown that neuroblasts that delaminate early and late not only position themselves at proximal and distal regions of the auditory ganglion but innervate the corresponding regions of the BP. This may imply that tonotopic identities of future auditory neurons are augured by time of delamination of precursors as they leave the PNSD of the OV. However, conclusive interpretations can only be drawn using perturbation studies, where neuroblasts delaminating at a specific time are prevented from leaving the epithelium or induced to delaminate at an earlier time.

### Encoding Time during Delamination

Our experiments show that differences in delamination timing specify the positional identity of neuroblasts in the inner ear ganglion. Moreover, this correlates with the innervation of both peripheral and central targets. This raises the question: How could time be encoded during the delamination of the inner ear neuroblasts? One example is seen in the segmentation clock during somitogenesis, where the formation time of each somite shows a temporal periodicity based on the cyclic waves of Hes accumulation and degradation (Palmeirim et al., 1997; Kageyama, 2022; Pourquie, 2022). In the developing telencephalon, the Notch effector, Hes1 has been shown to undergo oscillations in neural progenitor cells. These oscillations induce oscillations of the proneural gene Neurogenin2 and Notch ligand Delta-like1 (Dll1).

Oscillations maintain a neural progenitor state, while sustained expression promotes differentiation (Shimojo et al., 2008, 2011).

A similar process may occur in the inner ear. The cells of the otocyst divide through a process called interkinetic nuclear migration (INM), where the nuclei move to different levels of the epithelium depending on the cell cycle stage. Division occurs apically, and neuroblasts then move basally (Lorenzen et al., 2015). Similar to the neuroblasts of the epibranchial placodes, otic neuroblasts continue their basal migration, leaving the epithelium through breaches in the basal lamina (Begbie et al., 2002; Graham et al., 2007; Davies, 2011).

Oscillations of Notch effectors, combined with the oscillations during division, may specify neuroblast specification and subsequent delamination. Consistent with this is the regulation of Ngn1 by Notch signalling, and the salt and pepper expression of Ngn1, Dll1and Hes5 in the otocyst during the stages at which neurons are being generated (Adam et al., 1998).

Furthermore, blockade of Notch signalling using DAPT, leads to the absence of Hes5 expression, increase in the expression of Dll1 and a dramatic increase in delaminating neuroblasts as evidenced through NeuroD1 upregulation (Abello et al., 2007). While this will specify the pace of neurogenesis, the changing properties of neuroblasts are likely to result from other factors in the otocyst.

## ACKNOWLEDGEMENTS

This work was supported by the Department of Atomic Energy, Government of India, Project Identification No. RTI 4006, and grants from SERB, TIFR Infosys-Leading Edge Grant, the Royal National Institute for Deaf People. We acknowledge the support of the Central Imaging Facility at NCBS and to the Central Poultry Development Organization & Training Institute, Bengaluru.

---

**Extended Data to Figure 1.**
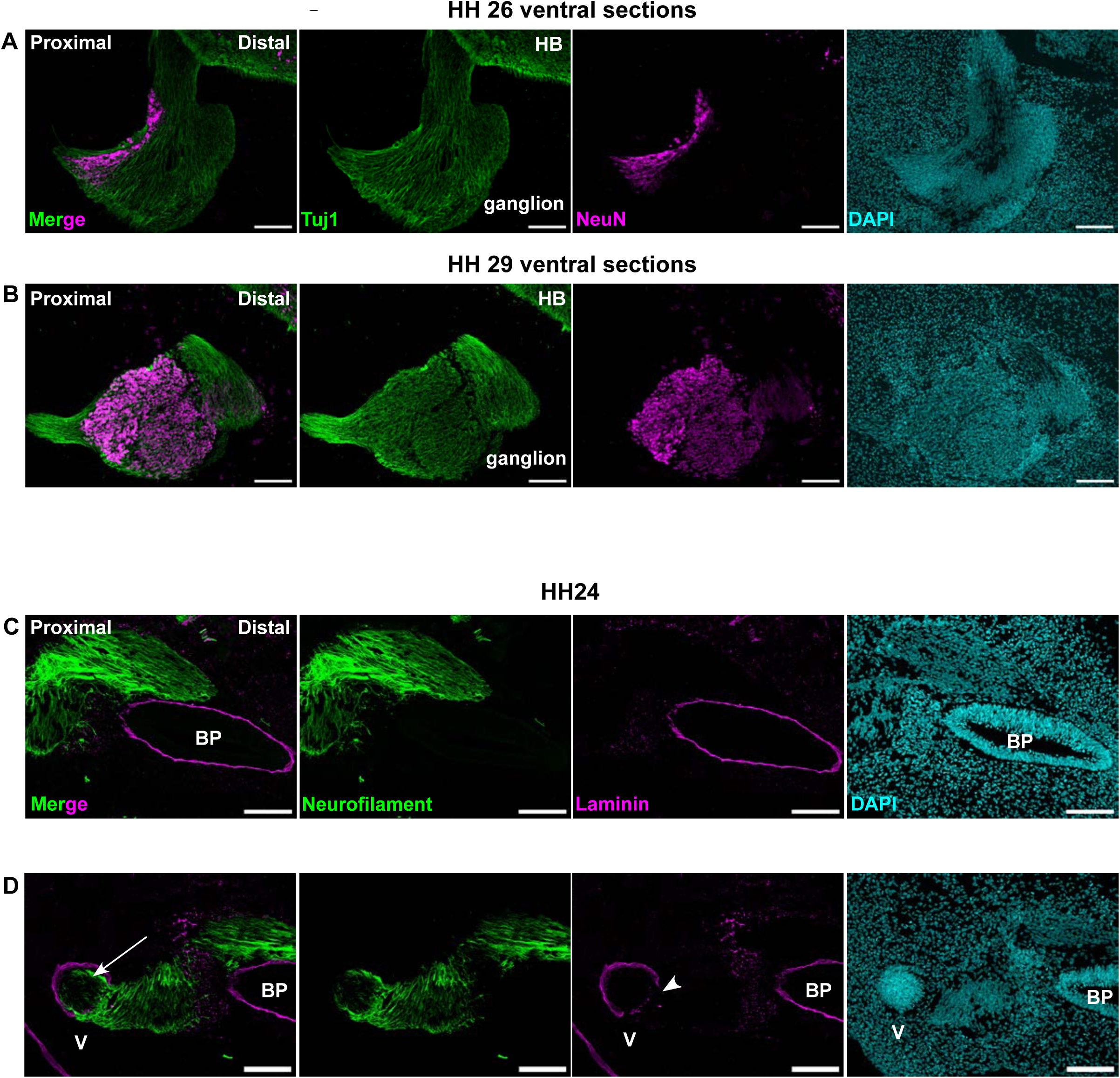
A - Ventral sections of inner ear ganglion of stage HH 26 embryo. B - Ventral sections of inner ear ganglion of stage HH 29 embryo. As seen in more dorsal sections, the post-mitotic neuronal differentiation marker NeuN can be seen spreading from proximal to distal regions of the ganglia as development proceeds. C - Sections of stage HH 24 chicken inner ear showing that neuronal fibres have not yet invaded the nascent BP. D - Sections of stage HH 24 chicken inner ear showing breaches in the laminin barrier of the vestibule (arrow head) and innervation of the same by neuronal fibres (arrow). Scale bar 100 μm

**Extended Data to Figure 2.**
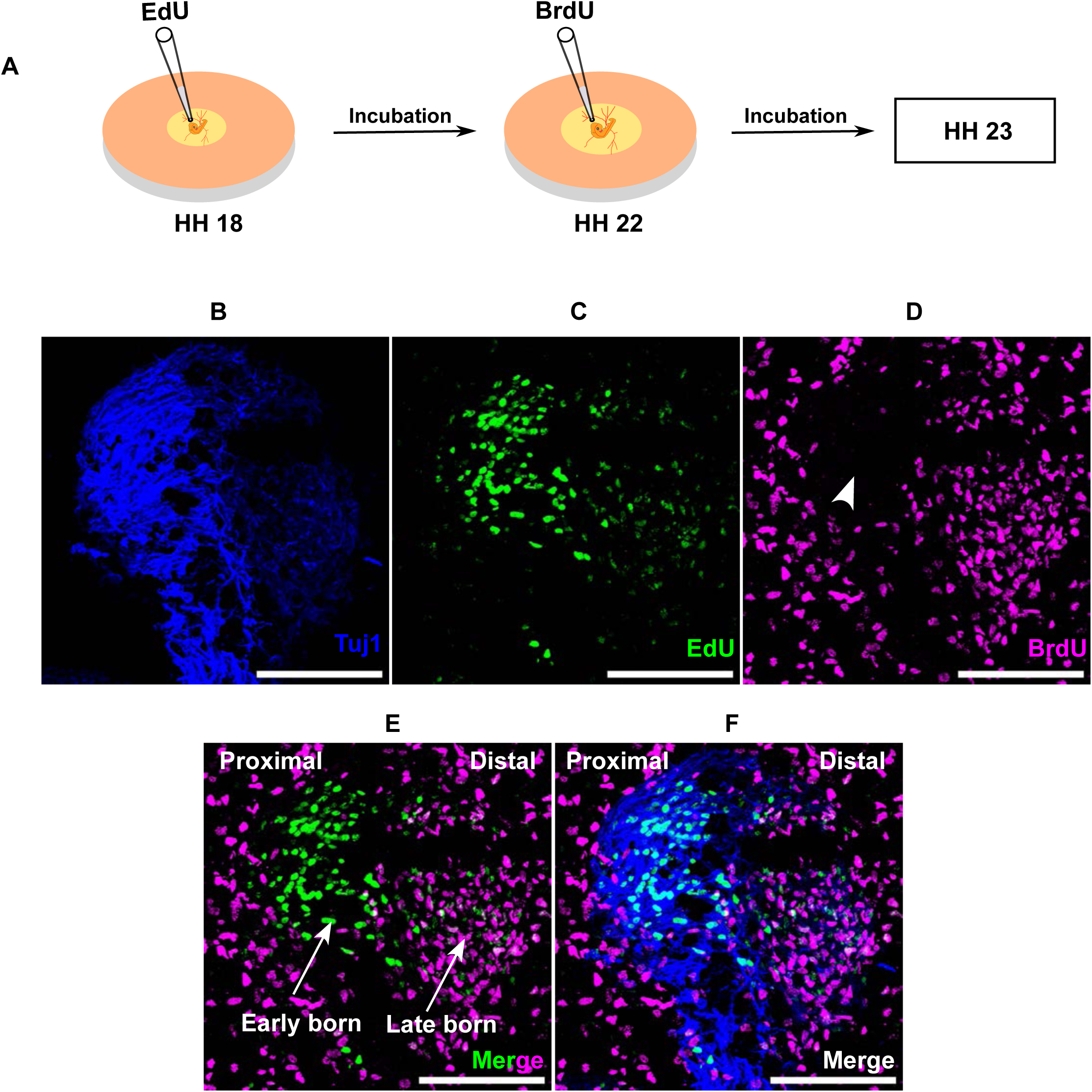
A - Schematic of protocol followed for sequential labelling of inner ear neurons born at different times. EdU labelling at stage HH 18 was considered early born while neurons labelled at stage HH 22 were called late born. Embryos could not be cultured beyond stage HH 23 with this protocol. B - Tuj1 labelling of inner ear ganglion. C - EdU labelling showing early born neurons. D - BrdU labelling showing late born neurons. Arrow head shows the absence of late born neurons in more proximal regions of the ganglion. E - Arrow heads show early born neurons are proximal and late born neurons are distal even at very early stages of development. F - Merge section. Scale bar 100 μm

**Extended Data to Figure 6:**
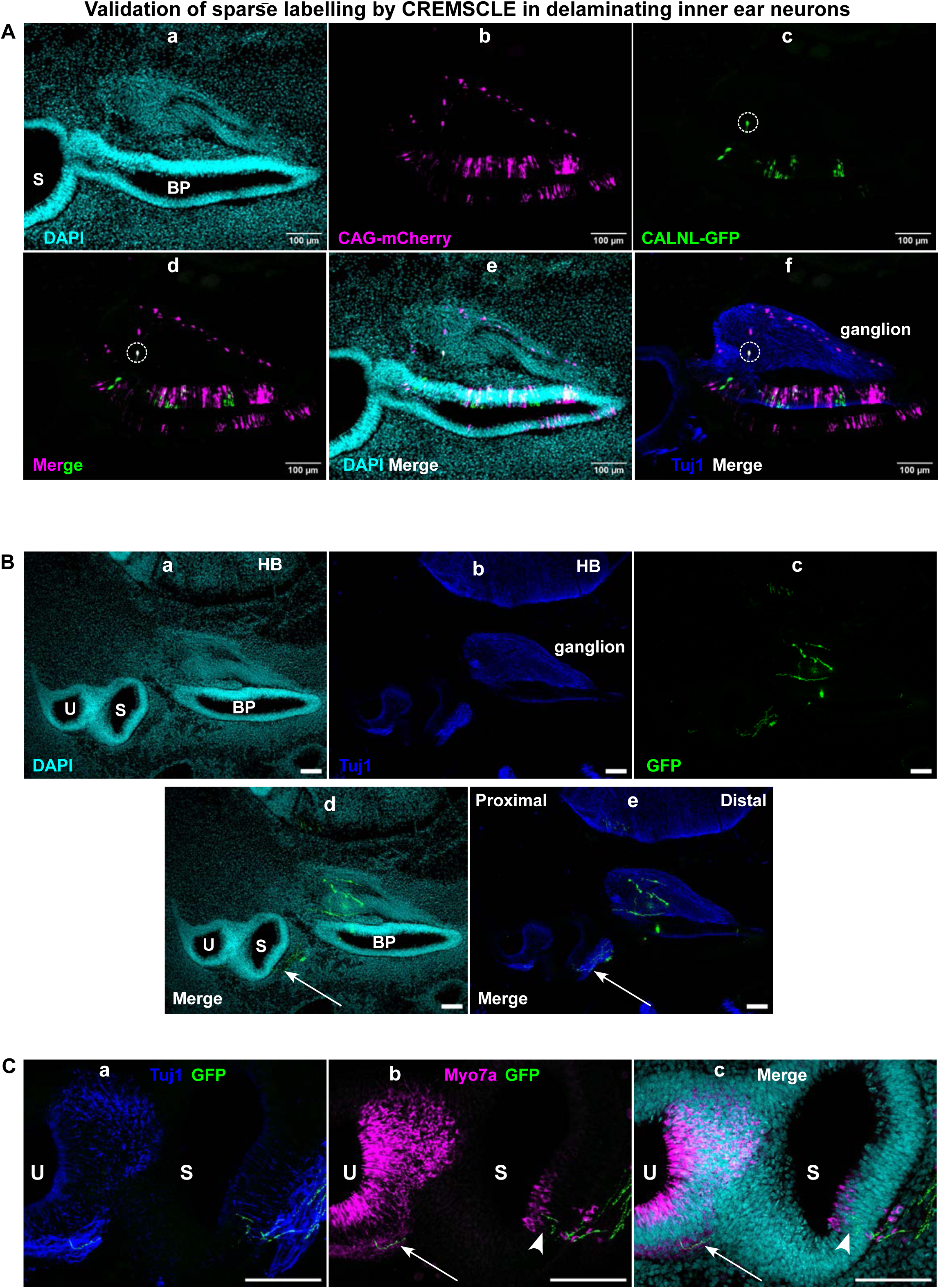
A-a to f - Validation of sparseness of GFP labelling by CREMESCLE. CALNL-GFP and CAG-Cre were co-electroporated with CAG-mCherry. More mcherry positive delaminated neurons are visible in the ganglion as compared to sparse labelled GFP positive neurons (dashed circle). B- a, b and c - Sections of stage HH 29 inner ear in which OV was electroporated with sparse GFP (CREMSCLE) at stage HH 18 for labelling early delaminating cells. B- d and e- Merge images in which arrows show fibres from early delaminating neurons innervating the saccule. (U- Utricle, S- Saccule, BP- Basilar papilla, HB- Hindbrain) C- a, b and c show the merged high magnification images of the vestibular region. Arrows in C- b and c show a fibre innervating the utricular region. Arrow heads in b and c show fibres of early delaminating neurons innervating the saccule and making contacts with the saccular hair cells. Scale bar 100 μm

## Notes

### Competing Interest Statement

The authors have declared no competing interest.

### Summary of Updates

Typos and problems with reference formatting have been updated

